# The progression of neurovascular features and chemokine signatures of the intervertebral disc with degeneration

**DOI:** 10.1101/2024.07.12.603182

**Authors:** Remy E. Walk, Kaitlyn S. Broz, Liufang Jing, Ryan P. Potter, Christian E. Gonzalez, Alec T. Beeve, Erica L. Scheller, Munish C. Gupta, Lori A. Setton, Simon Y. Tang

## Abstract

Inflammatory cytokine production and de novo neurovascularization have been identified in painful, degenerated intervertebral discs (IVDs). However, the temporal trajectories of these key pathoanatomical features, including the cascade of inflammatory chemokines and neo-vessel and neurite infiltration, and their associations with IVD degeneration, remain relatively unknown. Investigating this process in the caudal mouse IVD enables the opportunity to study the tissue-specific response without confounding inflammatory signaling from neighboring structures. Thus this study aims to define the progression of chemokine production and neurovascular invasion during the IVD degeneration initiated by injury in the caudal spine 3-month-old C57BL6/J mice. Forty-nine IVD-secreted chemokines and matrix metalloproteinases (MMPs) was measured using multiplex ELISA, and the intradiscal infiltrating vessels (endomucin) and nerves (protein-gene-product 9.5) was quantified in the tissue volume using immunohistochemistry. Injury provoked the increase secretion of IL6, CCL2, CCL12, CCL17, CCL20, CCL21, CCL22, CXCL2 and MMP2 proteins. The centrality and structure of inflammatory networks in IVDs evolved over the 12 post-injury weeks, highlighting distinct responses between the acute and chronic phases. Neurites propagated rapidly within 2-weeks post-injury and remained relatively constant until 12-weeks. Vascular vessel length was observed to peak at 4-weeks post-injury and it regressed by 12-weeks. These findings identified the temporal flux of inflammatory chemokines and pain-associated pathoanatomy in a model of IVD degeneration using the mouse caudal spine.

## 1. Introduction

Low back pain affects up to 85% of the population worldwide^1,2^, and intervertebral disc (IVD) degeneration is a significant contributing factor.^3^ The IVD is a cartilaginous soft tissue and is considered avascular and aneural.^4^ Sandwiched between vertebral bodies, the IVD provides resistance against compressive loads and shock absorbance for the axial skeleton. With aging and injury, the IVD degenerates with the compromised ability to perform these essential functions and ultimately leading to low back pain.^3^ In addition to the structural collapse and the depletion of proteoglycan-rich matrix, other hallmarks of the degenerating IVD may be culprits to chronic pain, including the production of inflammatory chemokines, expression of catabolic enzymes^5,6^, and the invasion of neurites and vessels.^7,8,9,10^ Chemokines canonically recruit immune cells, which in turn secrete more chemokines that further exacerbate the inflammatory state of the degenerating IVD^11^, and the immune cells can further accelerate the breakdown of the extracellular matrix.^12,13^ Chemokines produced by the IVD may also contribute to neuron and vessel propagation around and into the outer annulus fibrosus which may mediate of IVD degeneration associated low back pain.^14^ Further, chronic presence of these chemokines can sensitize nociceptive neurons to produce more pain signals.^58^

Animal models are a common tool for studying IVD degeneration. Injury such as via mechanical overloading^15–18^ or needle puncture^19–33^ are used to provoke degeneration of the IVD. Though the lumbar spine is more clinically relevant as a site of pain generation compared to the caudal spine, the surgical exposure required to access the lumbar IVD is traumatic, and the surrounding inflammation may confound the IVD-specific responses. Puncture injury to the caudal IVD does not require surgical exposure and can be implemented with relatively simple surgical exposure or radiographic guidance.^25^ Furthermore, the murine caudal spine consists of 27 intervertebral discs, compared to just 5 in the lumbar spine,^34^ and thus enable better control of inter-animal variability by allowing comparisons of IVDs subjected to different treatment conditions within the same mouse. Therefore, the caudal spine may be more experimentally efficient for investigating IVD-specific disease mechanisms.

To effectively leverage the advantages of the caudal model, it is crucial to define the progression of the inflammatory cascade and pain-related neurovascular features over time. Both neurites and vessels have been observed in aged mouse lumbar IVDs^35^ and in human degenerated IVDs^8^, but the time course of how the caudal IVD recapitulates these features is unclear. Therefore, the objective of this study is to define the temporal progression of neurites, vessels, and the local production of chemokines during injury-induced degeneration of the mouse caudal IVD.

## 2. Materials and Methods

### 2.1 Animal model

All animal procedures were performed with Washington University School of Medicine IACUC approval. Three-month-old C57BL6/J female mice (N = 35) were used in this study. They were housed under standard animal husbandry conditions (in a temperature-controlled [21lJ±lJ1°C] room with normal 12-h light/dark cycles). Bilateral puncture with 30G needle of caudal (Coccygeal - Co) intervertebral discs (IVD) was performed and adjacent IVDs were used as internal controls. Pre- and post-procedural X-ray (Faxitron UltraFocus 100) was used to locate the IVDs of interest to confirm puncture. Co4/5 and Co6/7 were injured with Co3/4 and Co7/8 acting as internal controls. A group of animals (n = 5) underwent a sham procedure to create a superficial injury where the only the skin and surrounding soft tissue was punctured without injury to the IVD. Longitudinal assessment of pain behavior and locomotive performance was performed on a subset of animals (Supplemental methods). The mice were euthanized at 2, 4 and 12 weeks (n = 9-10 per timepoint) after injury; all sham mice were taken out to 12 weeks. Paired control and injured IVDs from bilateral puncture mice were divided between OCT embedded histology (Co3/4 and Co4/5; n = 9-10 per timepoint), paraffin embedded histology (Co7/8 and Co6/7; n = 5 per timepoint) and organ culture (Co7/8 and Co6/7; n.= 4-5 per timepoint). Sham control and punctured IVDs were divided between immunohistochemistry (Co3/4 and Co4/5) and paraffin embedded histology (Co7/8 and Co6/7). The lumbar dorsal root ganglions were also extracted from a subset of animals and underwent staining for TRPV1 (Supplemental methods). All samples for histology were immediately fixed in 4% paraformaldehyde at time of sacrifice for 24-48 hours.

### 2.2 Paraffin embedded histology

Spinal segments with Co7/8 and Co6/7 (n = 5 per timepoint) were embedded in paraffin and 10 µm thick sagittal sections were stained with Safranin-O against FAST green. IVDs were graded using a standardized 35-point histopathology scale.^36^ Three independent blinded individuals graded all histological sections to consensus.

### 2.3 Quantification of Secreted Factors

Co7/8 and Co6/7 (n = 4-5 each per timepoint) functional spinal units (FSUs) were immediately placed in tissue culture media at time of sacrifice. Culture media consisted of 1:1 Dulbecco’s modified Eagle’s medium: Nutrient mixture F-12 (DMEM:F12) supplemented with 20% fetal bovine serum and 1% penicillin–streptomycin. FSUs were cultured for 6 days at 37°C and 5% CO_2_ with a complete media change after 3 days. Media that was collected on Day 6 was analyzed as described here. The chemokines were measured using the Luminex™ 200 system (Luminex) using two separate kits, a 32-plex and a 12-plex assays (MilliporeSigma) to detect a total of 44 markers. The 32-plex included Eotaxin (CCL11), granulocyte colony-stimulating factor (GCSF), granulocyte-macrophage colony-stimulating factor (GMCSF), IFN-γ, IL-1α, IL-1β, IL-2, IL-3, IL-4, IL-5, IL-6, IL-7, IL-9, IL-10, IL-12 (p40), IL-12 (p70), IL-13, IL-15, IL-17, IP-10 (CXCL10), KC (CXCL1), leukemia inhibitory factor (LIF), LIX (CXCL5), MCP-1 (CCL2), MCSF, MIG (CXCL9), MIP-1α (CCL3), MIP-1β (CCL4), MIP-2 (CXCL2), RANTES (CCL5), TNFα, and VEGFA. The 12-plex measured 6Ckine/Exodus2 (CCL21), Fractalkine (CX3CL1), IFN-β1, IL-11, IL-16, IL-20, MCP-5 (CCL12), MDC (CCL22), MIP-3α (CCL20), MIP-3β (CCL19), TARC (CCL17), and TIMP-1. Assay sensitivities of these markers range from 0.3–30.6 pg/mL. Matrix metalloproteases (MMPs) were quantified using a single 5-plex kit (MilliporeSigma). This kit measured MMP-2, MMP-3, MMP-8, proMMP-9 and MMP-12. Assay sensitivities of these markers range from 1.6 – 8.4 pg/mL. Individual analyte sensitivity values for all kits are available in the MilliporeSigma MILLIPLEX® MAP protocol.

### 2.4 Evaluation of intradiscal vascularization and innervation

The spinal segments including Co3/4 and Co4/5 IVDs (n = 9-10 per timepoint) were embedded in OCT and sectioned along the sagittal plane at a 50 µm thickness. Frozen sections were stained with anti-protein gene product 9.5 (PGP9.5) and anti-endomucin (EMCN) against DAPI. PGP9.5 is a neuronal marker for sensory and autonomic nerve fibers and EMCN is an endothelial cell marker. Visualization of PGP9.5 and EMCN was achieved with Alexa Fluor 488 (green) and Alexa Fluor 647 (red) antibodies, respectively.

Three-dimensional image stacks were obtained via confocal fluorescence microscopy (DMi8, Leica Microsystems) and a maximum intensity image of each 50 μm section was generated for analysis. Nerves and vessels were semi-automatically traced using ImageJ 2.3.0 SNT plugin.^37^ Individual structure lengths were tabulated and total neurite and vessel length was calculated including both the posterior and anterior sides of the IVD. The outer annulus fibrosus and immediately adjacent tissues were included as the region of interest (ROI) for quantification.

### 2.5 Cytokine Network Analysis

To further characterize the temporal variation in inflammatory signaling, networks of cytokine interactions were constructed and analyzed using a custom MATLAB (Version: 9.13.0.2080170 R2022b) script. Networks were generated by calculating a Pearson correlation matrix for each timepoint by ratioing each analyte’s concentration between injured and uninjured discs for each animal. To focus only on strong protein correlations, a threshold (|r| > 0.7) was applied to the correlation matrices and self-loops were removed. The filtered matrices were used to create undirected graphs, with nodes representing cytokines and edges representing significant interactions. Eigenvector centrality and betweenness centrality were computed to determine important cytokines for each timepoint network. High-ranking cytokines shared between timepoints and unique to each timepoint were identified. Additionally, key network characteristics were extracted to understand the structure and function of the cytokine networks. The path length was computed as an average of the shortest finite paths between all pairs of nodes; modularity was computed using the Louvain community algorithm (Blondel et al., 2008). The Jaccard index calculated for all pairs of networks with 2-hop reachability matrices to allow for quantifying similarity between networks with a slight tolerance for indirect edge comparisons. Finally, networks were visualized using force-directed layouts with nodes colored by eigenvector centrality and sized by betweenness centrality.

### 2.6 Statistics

A paired two-way ANOVA was used to test for an effect of injury and week post-injury between the experimental and control segments, at a significance level of 0.05 with a post hoc Tukey HSD (Prism 10.2.2, GraphPad). A paired t-test was used to test for an effect of the superficial injury on experimental levels versus control segments in the sham-injured group only (12 weeks).

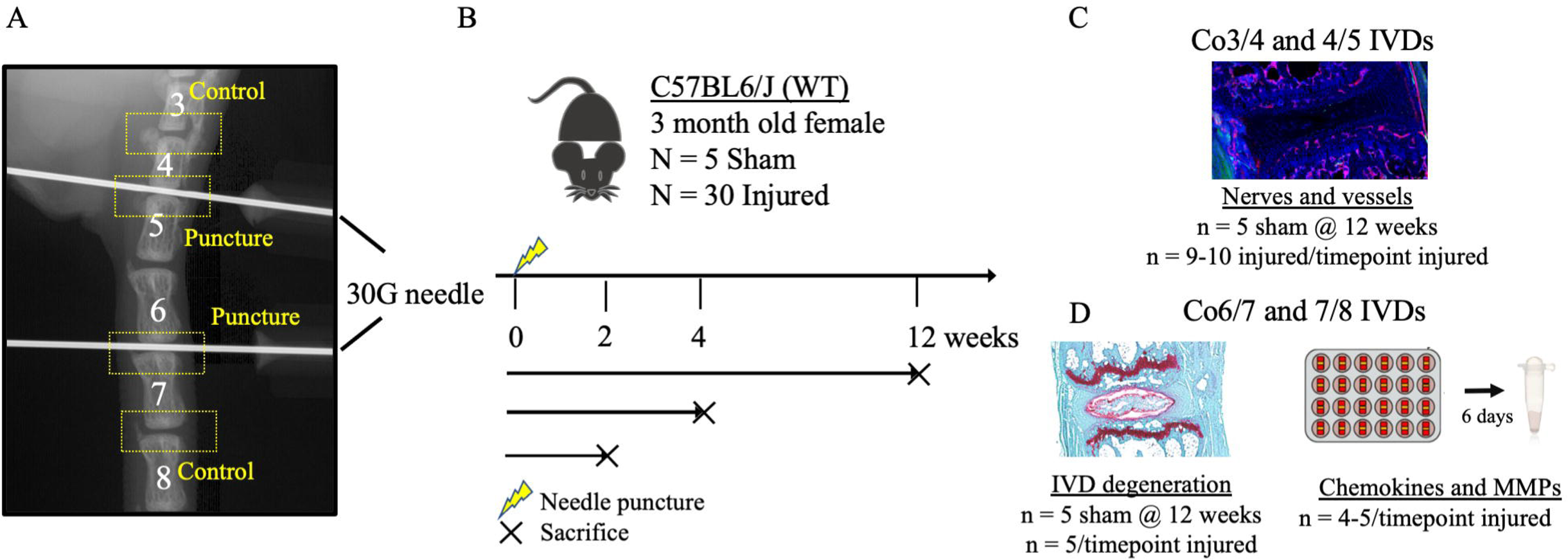

## Results

### 3.1 Direct injury to the intervertebral disc causes rapid and sustained degeneration

Bilateral puncture of the caudal intervertebral disc (IVD) resulted in mild to severe IVD degeneration (Fig 2A). Complete collapse of the IVD was observed in the most severe cases. Co6/7 and Co7/8 IVDs from sham and bilateral puncture mice were graded on a histopathologic scale for IVD degeneration and total IVD grade was significantly increased in punctured IVDs compared to internal controls of injured (p<0.05, ANOVA) but no differences were detected in these levels in the sham mice (p=0.1, t-test; Fig. 2B). Multiple compartments of the IVD showed degenerative changes, including the nucleus pulposus, annulus fibrosus and the interfaces at all timepoints after injury while cartilaginous endplates were only significantly degenerated at 12 weeks following injury (Fig. 2C). No effect of injury was observed in any pain behavior or locomotive assessments {Supplemental}.

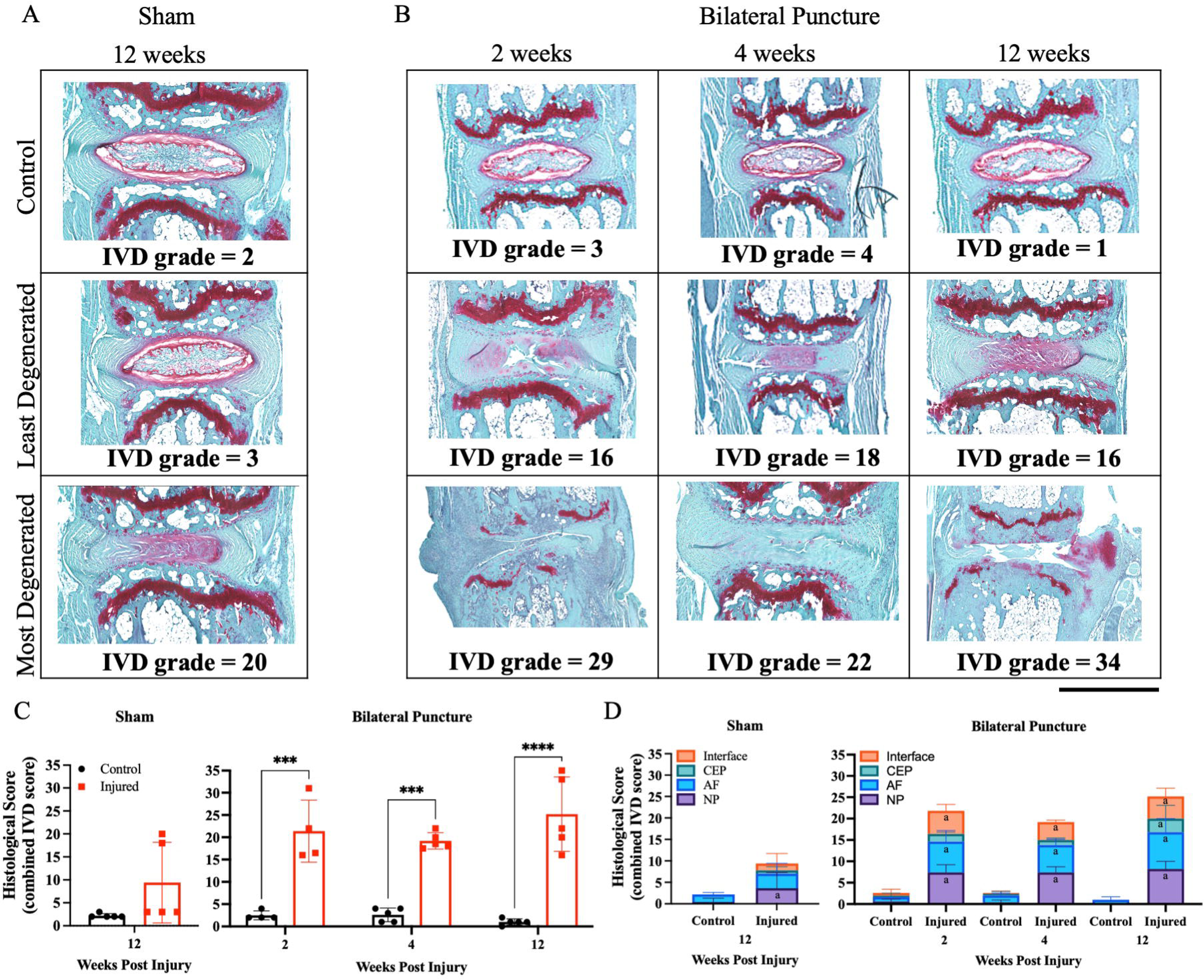

### 3.2 Chemokine production peaks at 2 weeks after injury

Forty-four distinct chemokines and five MMPs were measured from the culture media of control and punctured functional spinal units from injured mice. An effect of injury was seen in both pro-inflammatory chemokines (IL6 and TNFα) and immune cell recruitment chemokines (CCL4, CCL12, CCL17, CCL20, CCL21, CCL22 and CXCL2) (Fig 3). TNFα and IL1β are pivotal inflammatory chemokines in IVD degeneration, and in this experimental model we only see increased TNFα expression.^38,39^ The greatest difference between injured and control IVDs in chemokine production occurred 2 weeks after injury where significantly higher expression of CCL12, CCL17, CCL20, CCL22 and TNFα with all but CCL20 returning to control levels by 4 weeks post injury and CCL20 by 12 weeks post injury (Fig 3A-E). CCL21 was elevated at 12 weeks after injury (Fig 3G). MMP-2 was detected as being affected by injury with the peak at 12 weeks post injury (Fig. 3J). Approximately x chemokines were not detectable at any of the time points, while Y chemokines were not different over time (Supplemental Table 1).

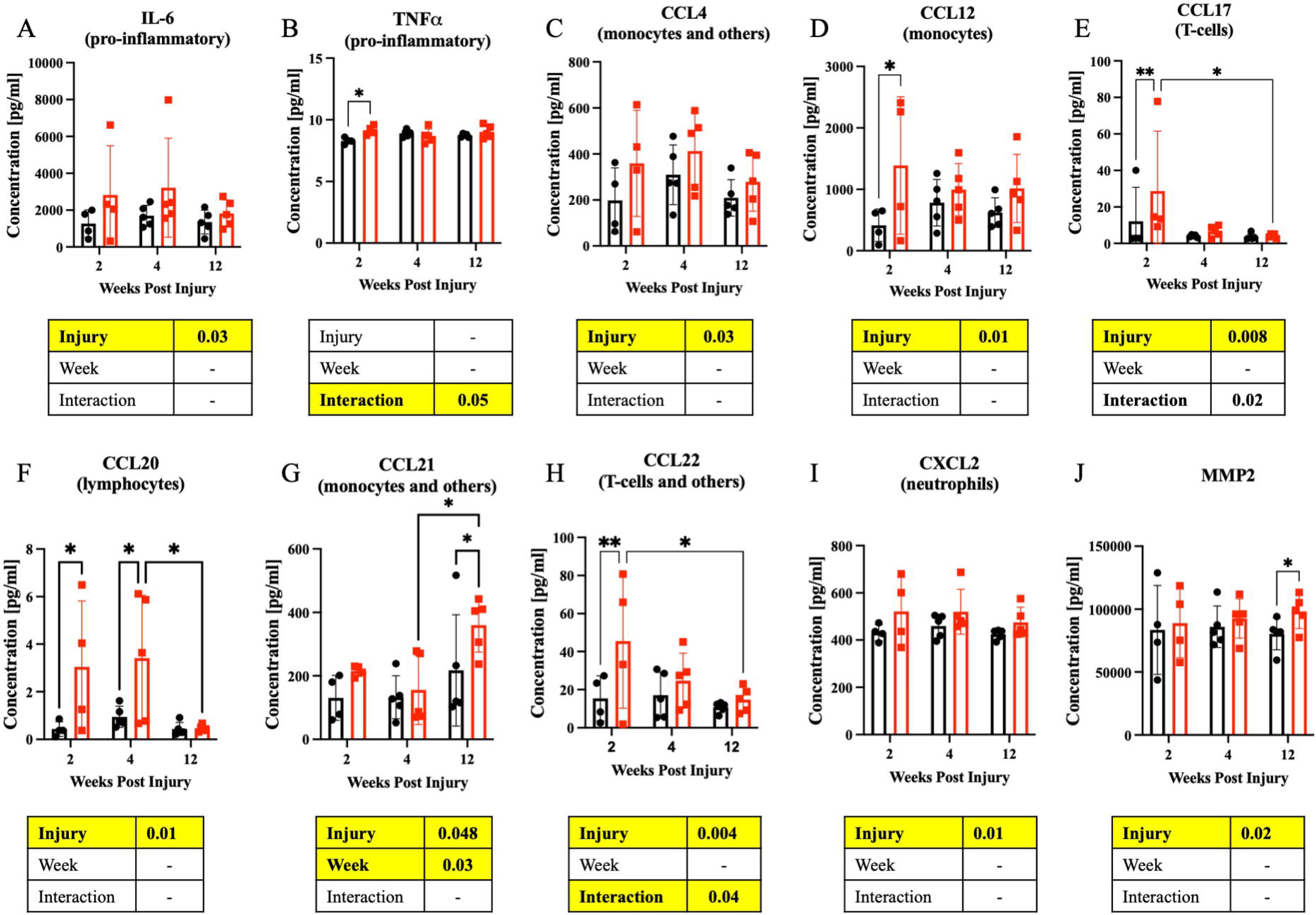

### 3.3 Innervation and vascularization propagate at different temporal trajectories

PGP9.5+ neurite and EMCN+ vasculature structures were manually segmented on a maximum projection image (Fig 4A). The region of interest (ROI) contained the anterior and posterior outer annulus fibrosus and surrounding tissue. High magnification ROIs show innervation and vascularization that colocalize in these areas (Fig 4B). Nerve and vessel structures were semi-automatically traced and lengths were tabulated in each IVD.

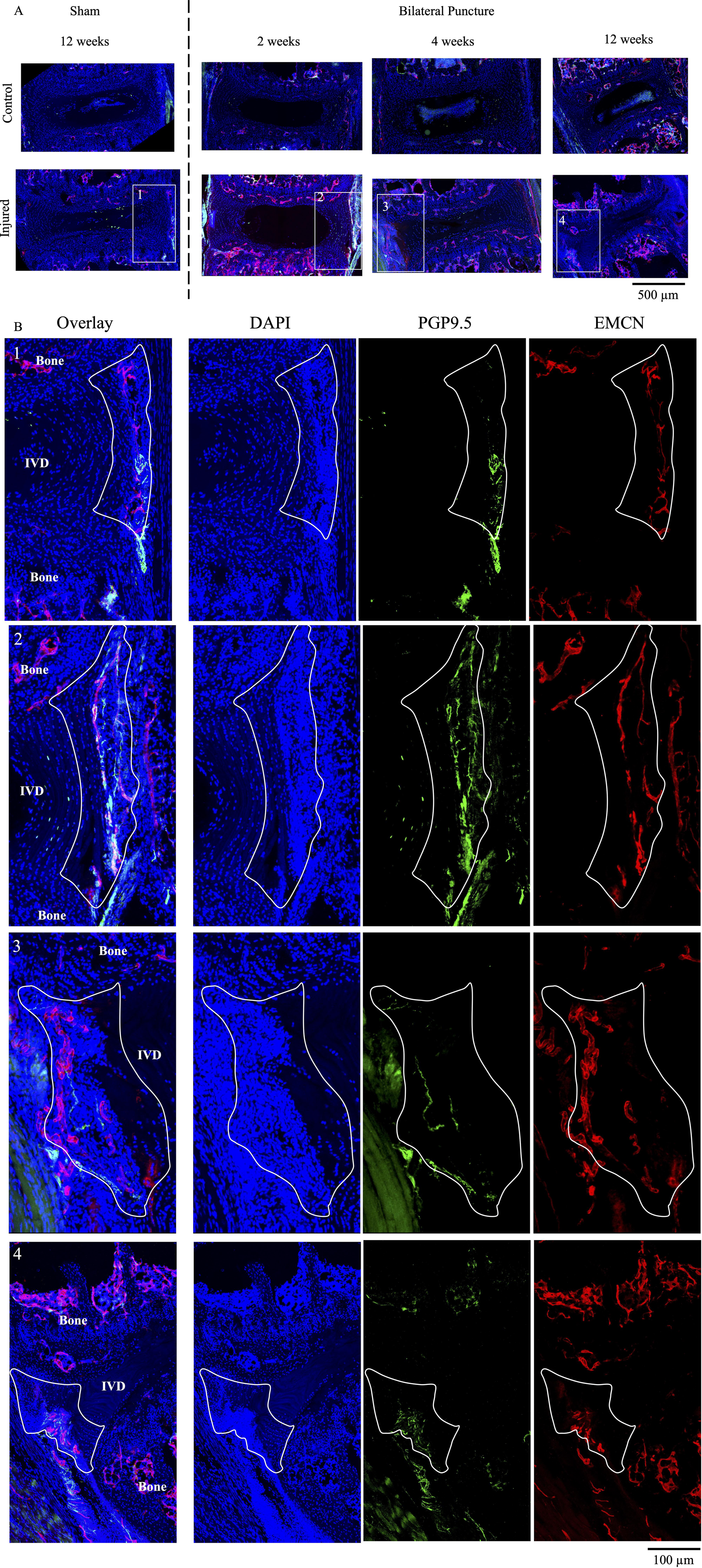

Sham mice showed negligible amounts of innervation and vascularization within their IVDs. Total length of each feature in each IVD was measured and an increased presence of both structures was observed as early as 2 weeks after injury (Fig 5A-B). PGP9.5+ neurite structures are observed 2 weeks following injury and remain consistently increased through the 12 week period; in comparison, EMCN+ vessels peak at 4 weeks and appear to recede by 12 weeks after injury. Violin plots of punctured IVDs from injured mice show the tabulation of individual nerve and vessel lengths that were measured with the total number of structures written above the plot (Fig 5C-D). The distribution of nerves remains consistent through all 12 weeks while the number of vessels in the 150-300 µm range is dramatically reduced at 12 weeks compared to 4 weeks post injury.

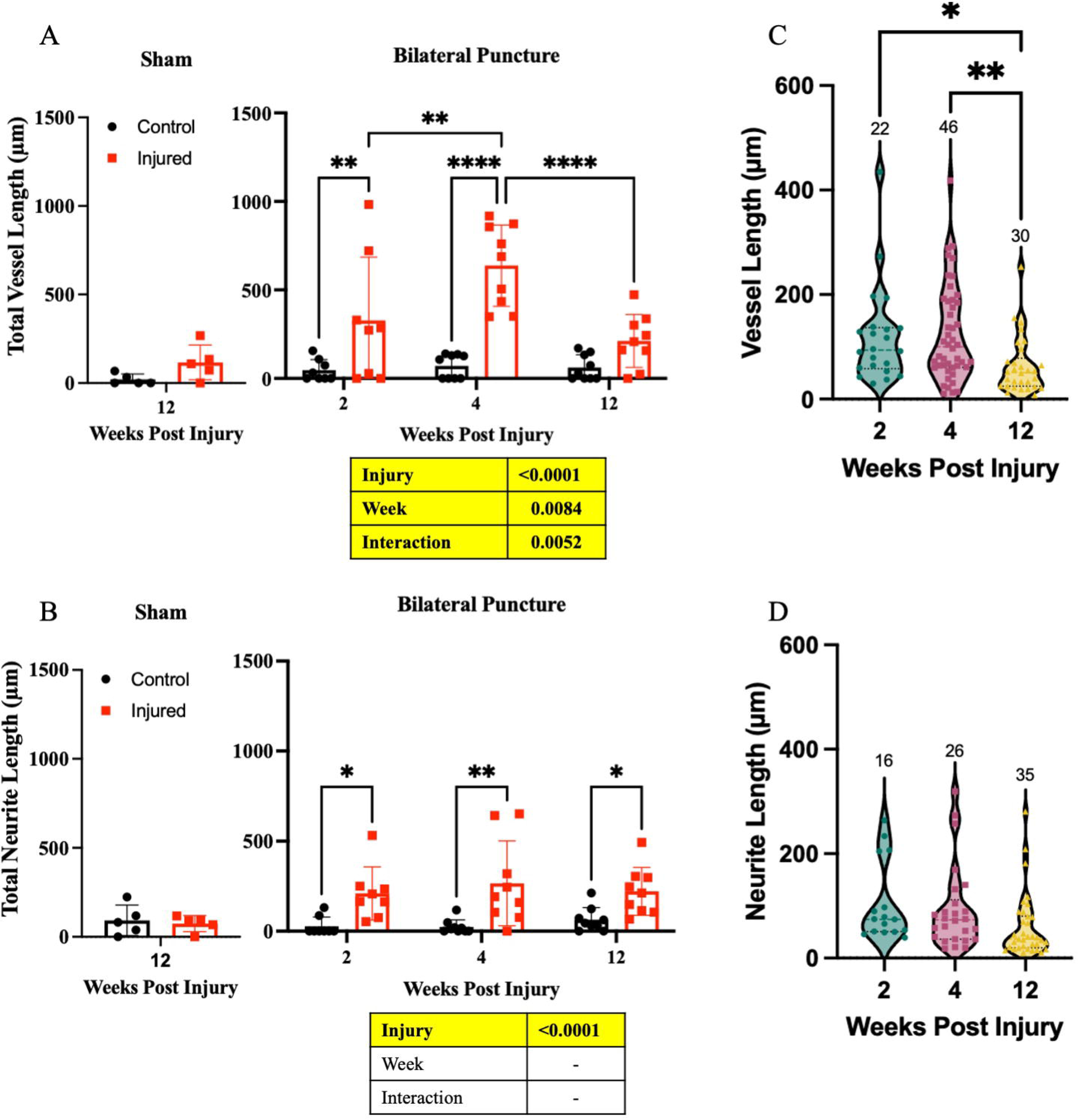

3.4

There were several novel findings in identifying influential cytokines in the networks across the 3 timepoints in this study. At 2 weeks post injury, pro-inflammatory (IL-4) and immune-cell recruiting (IFNγ, CCL2, CCL5) cytokines are high ranking in centrality. By week 4, these are no longer highly central, but IFNB1, IL12p70, and CXCL1 (which are immune-cell recruiting cytokines also highly central in the week-2 network) are still high-ranking nodes in centrality measures. Additional chemotactic and immune-cell regulating cytokines (CX3CL1, IL-16, and LIF) are highly influential in the network at week 4. In moving to week 12, a unique set of cytokines and pleiotropic factors (IL9, CXCL2, CXCL9, CCL17, CCL20, and VEGF) are central to network activity. Throughout all time points, IL-11 and CCL4 were consistently highly ranked in both centrality measures. In analyzing the network characteristics, modularity greatly increases as time progresses after injury (0.269, 0.368, 0.466 for week 2, 4, 12 respectively). This finding is corroborated by the increase in path length as time progresses (1.41, 1.69, and 1.80), wherein regulatory relationships become more distinct and linear while being less interactive and autoregulatory at later timepoints. In comparing network intersections, week 2 and week 4 are the most similar (0.327 Jaccard Index) while week 2 vs week 12 and week 4 vs week 12 are equally dissimilar (0.238 and 0.237 Jaccard Index respectively).

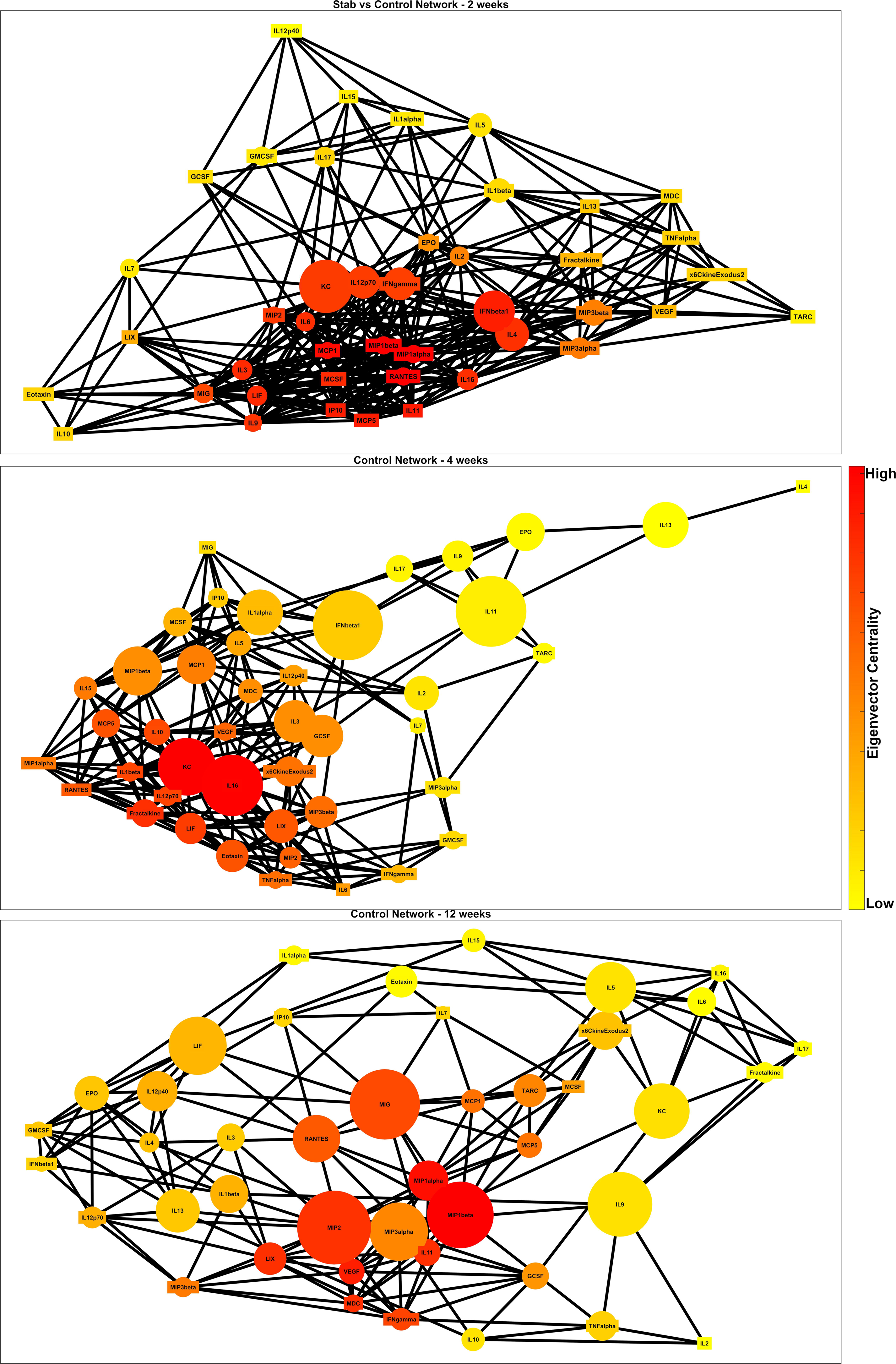

## Discussion

IVD injury models are a commonly utilized tool for studying the progression of IVD degeneration. In contrast to the lumbar spine, where degeneration is known to evoke changes in pain behavior^24,40^, we did not observe any behavioral changes following injury in the caudal spine (Fig. S1). Thus, the caudal IVD injury model is best suited to evaluate IVD-specific responses during degeneration. The advantage of a non-invasive surgical approach and access to multiple levels promotes the reduction in number of research animals used in accordance to the 3R principle.^41^ The use of the caudal spine also minimizes the interference due to the disruption and inflammation of surrounding tissues compared to the complex surgical access to the lumbar spine. While there have been extensive studies showing the structural and compositional degenerative changes following caudal puncture^25,29,30,42,43^, limited data exists on additional aspects of IVD degeneration including innervation^27,28^ and vascularization and chemokine secretion from the explanted IVD.^44^ Our results here show that the caudal IVD produces a significant amount of diverse chemokines, and it is susceptible to developing pain-associated features after injury.

Bilateral puncture of the caudal IVD resulted in quick and sustained IVD degeneration up to 12 weeks post-injury. Both proinflammatory (IL6 and TNFα) and chemokines (CCL4, CCL12, CCL17, CCL20, CCL21, CCL22 and CXCL2) were elevated with injury, with the highest expression of a subset of chemokines compared to controls at 2 weeks following injury. These chemokines canonically recruit monocytes, T-cells, and lymphocytes.^45^ Yet chemokines are known to be pleiotropic and have been associated with additional functions such as IVD degeneration, pain, neurite growth and angiogenesis.^46^ For example, CCL4 has been shown to be elevated in degenerated human IVDs and associated with pain behavioral changes in a rat model of IVD degeneration.^6,47,48^ TNFα injected into rat lumbar IVDs has been shown to enhance pain behavior changes, possibly through irritation of nerve endings.^33^ CCL17 and CCL21 induced dorsal root ganglion (DRG) axonal growth ^49,50^ and CXCL2 is a known mediator of angiogenesis.^51^ CCL17 and CCL22 through the receptor CCR4 were indicated to play a role in pain development and CCL22 was able to activate neurons and increase neuron excitability.^52^ Chemokines production of the IVD following injury may help provide further insights into the pathoanatomy of innervation and vascularization as well as provide possible pathways for IVD degeneration associated low back pain.

Correlative network analysis of the cytokine production revealed several key factors in both the acute and chronic phases of injury. CCL4, a factor significantly upregulated in the IVD within this injury model, is highly central in the cytokine network at all time points. In contrast, factors like CXCL2, CCL17, and CCL20 are also upregulated during injury but are only highly central at the latest timepoint of 12 weeks. This highlights two distinct regimes of chronic inflammatory degeneration within the IVD. Initially, factors like CCL4 may serve as directors early on and remain notably influential within the cytokine network all the way till the chronic timepoint of 12 weeks. In the end, however, there is a latent inflammatory network change in which CXCL2, CCL17, and CCL20 become prominent only during this final timepoint. This suggests that CCL4 may be a target for early intervention given its consistent influence on the expression of other cytokines, elevated expression during injury, and its suggested role in IVD degeneration and pain development previously mentioned. On the other hand, CXCL2, CCL17, and CCL20 may be more appropriate as targets of late-stage intervention in chronic painful degeneration of the disc, given their aforementioned roles in angiogenesis, pain development, immune cell recruitment.

Network comparison revealed that both modularity and path length increase greatly with time after injury. This suggests that earlier timepoints are characterized by a broad variety of inflammatory pathways functioning in parallel (a notion supported by the peak upregulation of cytokines at 2 weeks post injury). Contrarily, later timepoints are characterized by a more specific and linear set of chronic cytokine relationships. This is further supported by the network similarity scores, with week 12 being highly dissimilar from week 2 and week 4. This ultimately suggests that the microenvironment of the injured IVD switches from promoting acute inflammation to promoting chronic inflammation between week 4 and week 12, which could serve as the critical window for the mechanistic underpinnings of chronic inflammatory signaling to develop.

Innervation of the IVD may be the potentiator of low back pain observed with lumbar puncture models and this feature is recapitulated here in the caudal spine. Studies have previously illustrated innervation of the IVD following injury with detection of PGP9.5+ or CGRP+ staining injured IVDs, but without any quantification of the structures.^19,26,28^ Further, the coincidence of vascularization with neo-innervation has been previously observed but the time course of vessel propagation into the IVD following injury has not been documented. To overcome these limitations, we semi-automatically traced neurite and vascular structures on maximum projection images of PGP9.5 and EMCN stained thick sections.^37^ This allowed for the tabulation of neurites and vessels present in the region of interest and their lengths for comparison. We observed a time-dependent vessel infiltration of the outer annulus fibrosus and surrounding tissue. In contrast, neurites quickly infiltrated within two weeks of injury and remained at similar levels in the subsequent time post injury. It is likely that the penetration of the IVD by vessels would be considered prerequisite to infiltration by circulating cells, including monocytes and other immune cells that might be responsible for secretion of the chemokines.

Behavioral assays can be used following lumbar puncture to quantify pain.^20,23,24,26^ A caveat of the caudal puncture model is that it does not produce axial low back pain as it does not endure the axial torso loadings. Correspondingly, we observed no differences in behavioral measures between sham and bilateral puncture mice. Although not measured here, there may have been localized measures of pain including sensitization of the tail to mechanical and thermal stimuli (e.g., Hargreave’s test, tail-flick).^28,53^ Another possible surrogate of pain-related change is to quantify molecular expression of neurotrophic factors in the innervating lumbar dorsal root ganglia (DRG). The DRG has been linked to chronic pain, and the increase in the expression of pain-related neuropeptides as well as neuronal excitability may be the mediators of discogenic pain.^54^ Ongoing work utilizes immunohistochemical staining for altered presence of neurotrophic factors in lumbar DRGs that have a demonstrated role in mediating pain transmission in the spine such as transient receptor potential cation channel subfamily V member 1 (TRPV1).

The method for quantifying innervation and vascularization of the IVD enabled measuring these features with greater fidelity. Protein analysis the IVD secreted chemokines revealed potential molecular mediators of IVD degeneration, innervation and vascularization with relevance to generation of inflammation and pain. Many of the secreted chemokines found to be elevated may be associated with increased presence of infiltrating monocytes that may include macrophages, B-cells or T-cells.^55,56^ Not surprisingly, the key angiogenic factor, VEGFA, was not elevated at any of the measured timepoints. VEGFA is critically expressed early following tissue repair promote early angiogenesis, ^55^ and by two-weeks following injury VEGFA has already exerted its effects as evidenced by robust vessel formation. In this study, we intentionally measured locally produced chemokines which will remove the effects of systemic changes in the animal. Recent work shows that the chronic NFκB activation in the caudal IVD produces a secretome that promote macrophage migration.^57^ Our data here confirm that a degeneration-causing injury will upregulate a plethora of chemokines that will likely recruit multiple immune cell types^55^, concomitant with increasing neurovascular features. Future studies quantifying the presence of these immune cells would advance our understanding of a role for local versus systemic changes in modulating chemokine secretion, as well as key factors that govern the infiltration of these pain-associated features.

## Acknowledgements

We gratefully acknowledge the support of the Washington University Musculoskeletal Research Center. Multiplex Luminex cytokine panels were analyzed by Eve Technologies Corp.

