## Supplementary material for "The progression of neurovascular features and chemokine signatures of the intervertebral disc with degeneration": {Supplemental}

**1. Supplemental Methods**

1.1 *Behavioral testing*

A suite of behavioral assays was performed on the sham (n = 5) and 12-week cohort of injured mice (n = 10). Open field^1–3^, electronic von Frey (E-von Frey)^1,4^, grip strength^5^, rotarod^1–3^, inverted wire hang^6^, and hot/cold plate^7,8^ (Bioseb, France) were conducted prior to and 2, 7 and 12 weeks after injury. Open field monitors passive activity over an hour period with positions and speed measured every 5 minutes. Mice are placed in an open field arena (40 cm x 40 cm, with 30 cm high walls) equipped with an automated tracking system (Omnitech Electronics, Columbus, OH). E-von Frey tests sensitivity to mechanical stimuli of the hind paws; a probe attached to a load sensor is applied to the plantar surface of the hind paw until a response such as withdrawal is observed. The maximum load applied is recorded and the median value of three tests is used for analysis. Average of left and right hind paw is reported. Grip strength testing operates by having the mouse positioned parallel to the table and allowed to grab a pull bar with the fore paws. The mouse is then pulled backwards by the tail until fore paws are released with the maximum load reported. Average of three tests is used for analysis. Rotarod assesses the mouses motor coordination and balance. Mice are placed on an constantly accelerating rod (4-40 rpm over 2 minutes) and time spent on the bar is recorded. The maximum time of five tests is used for analysis. Prior to the official tests, mice are trained to walk on the rod at 4 rpm for 2 minutes. Mice were excluded if unable to meet this baseline capability. Wire hang is used to measure neuromuscular strength and endurance. Mice are place on a wire mesh (2 mm diameter) which is then inverted to 180°. Time until mouse falls, up to 2 minutes, is recorded. Maximum time of the three trials is used for analysis. Cold and hot plate are used to assess sensitivity to thermal stimuli. For both assays, mice are placed on an enclosed plate heated to 55±0.5°C or 5±0.5°C for hot and cold plate, respectively. Mice are observed until a response such as licking the paws is observed up to 15 second for hot plate and 50 seconds for cold plate. An hour of acclimation is performed before each assay.

1.2 *Dorsal root ganglion (DRG) immunofluorescence*

The lumbar spines were immediately fixed in 4% PFA for 24-48 hours at time of euthanasia. L4 and L5 DRGs were isolated, incubated in 30% sucrose and embedded in OCT before being sectioned transversely at a 7 µm thickness. DRGs were stained with transient receptor potential cation channel subfamily V member 1 (TRPV1; abcam ab6166) with Alexa Fluor 488 secondary against DAPI. A single section per mouse was imaged on a fluorescence confocal microscope (Leica SPE, Leica Microsystems). DRG neurons were manually traced using ImageJ and total number and TRPV1+ neurons were counted. Data is reported as percent TRPV1+ neurons of all identified neurons.

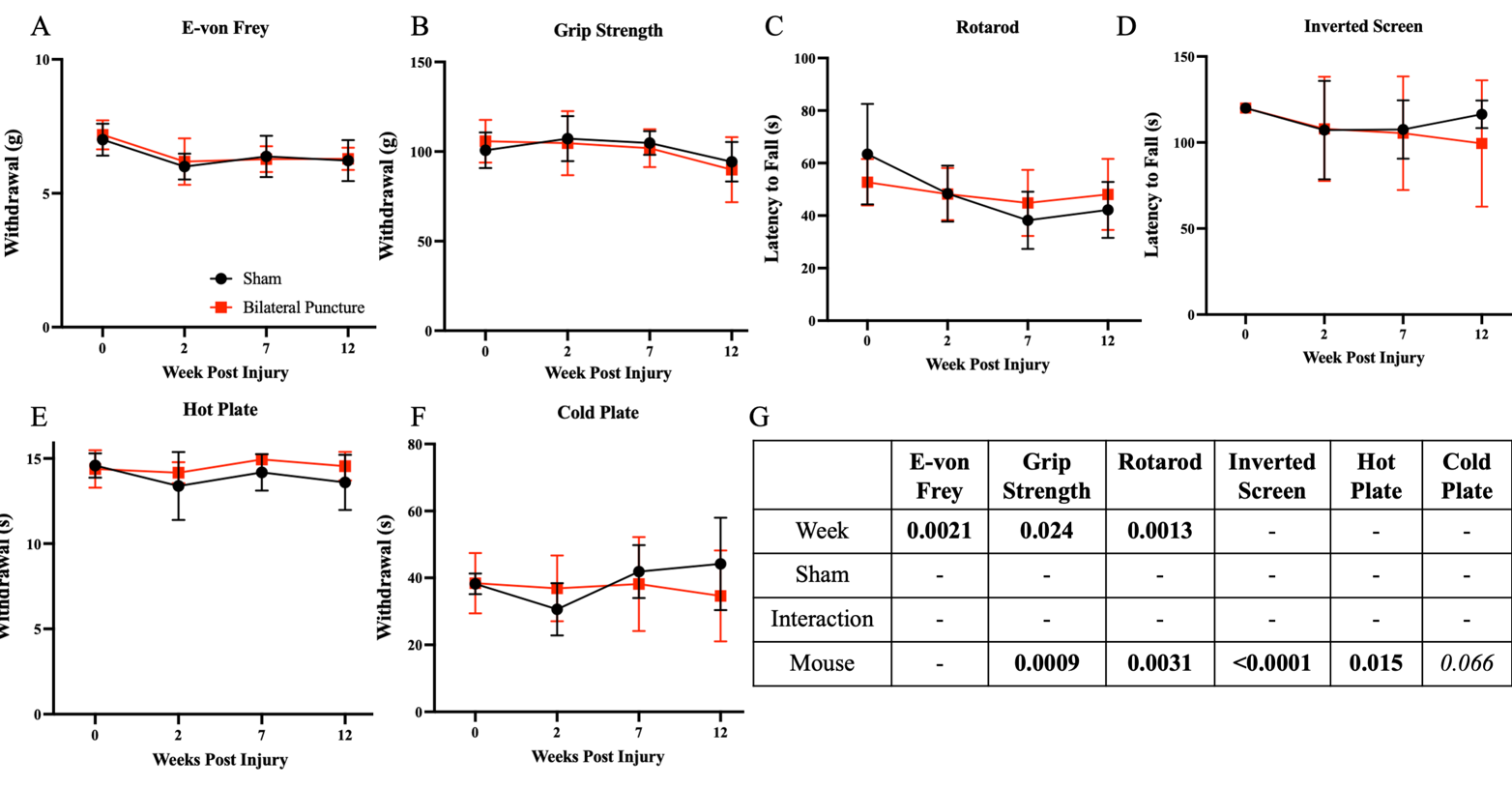
**2. Supplemental Data**

Figure S1: Behavioral assessment. No differences between sham and injured mice were detected. (A) E-von Frey, (B) grip strength and (C) rotarod had a significant effect of time. E-von Frey is reported as the average of left and right hind paw median withdrawal threshold. In addition, grip strength and rotarod as well as (D) inverted screen, (E) hot plate and (F) cold plate showed a significant effect of mouse. (G) Paired two-way ANOVA statistics shown.

Figure S2: Dorsal root ganglion (DRG) immunohistochemistry. Preliminary data on TRPV1 expression in lower lumbar DRGs following caudal puncture. Total neurons and TRPV1+ neurons were counted on stained sections against DAPI. (D) Percent TRPV1+ neurons is quantified. We see a larger average TRPV1+ neurons at 12 weeks compared to Sham and 2 weeks. Representative (B) Sham and (C) 2 and (D) 12 weeks post injury TRPV1 (green) stained against DAPI (blue).

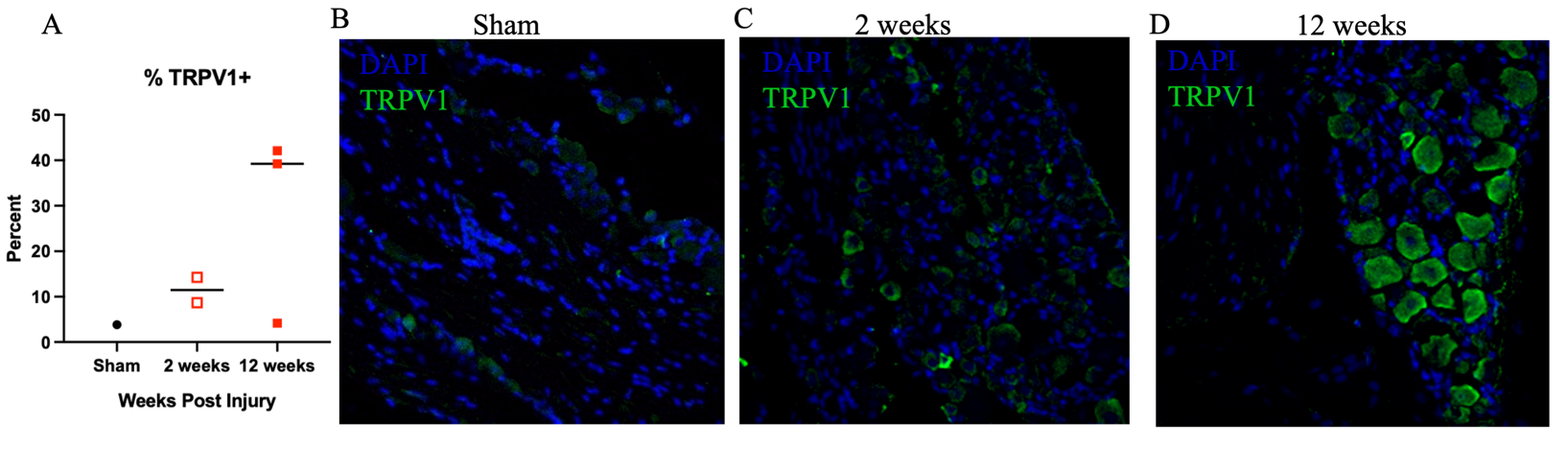

Supplement Table 1: Chemokines and Matrix Metalloproteinases

|  |  | Week Post Injury (Mean ± SD [pg/ml]) | | | P-values | | |
| --- | --- | --- | --- | --- | --- | --- | --- |
|  |  | 2 weeks | 4 weeks | 12 weeks | Interaction | Week | Injury |
| CC11 | Control | 102.14±50.48 | 173.23±66.10 | 187.67±61.40 | 0.90 | 0.10 | 0.72 |
|  | Injured | 117.81±74.98 | 168.85±32.77 | 196.27±79.96 |  |  |  |
| GCSF | Control | 67.60 ± 27.89 | 144.34 ± 75.67 | 111.88 ± 44.52 | 0.97 | 0.22 | 0.54 |
|  | Injured | 80.83 ± 51.04 | 150.21 ± 95.44 | 118.83 ± 53.69 |  |  |  |
| GMCSF | Control | 28.51 ± 1.26 | 28.63 ± 1.44 | 30.08 ± 1.06 | 0.44 | 0.34 | 0.94 |
|  | Injured | 29.60 ± 2.02 | 28.68 ± 2.10 | 29.09 ± 0.77 |  |  |  |
| IFN-γ | Control | 9.34 ± 0.36 | 9.32 ± 0.87 | 9.18 ± 0.58 | 0.43 | 0.67 | 0.71 |
|  | Injured | 8.88 ± 0.90 | 9.55 ± 0.65 | 9.17 ± 0.60 |  |  |  |
| IL-1α | Control | 113. 86 ± 6.05 | 118.35 ± 6.43 | 117.68 ± 5.84 | 0.23 | 0.69 | 0.25 |
|  | Injured | 118.74 ± 2.84 | 116.11 ± 4.79 | 121.30 ± 10.22 |  |  |  |
| IL-1β | Control | 25.98 ± 0.15 | 26.11 ± 0.75 | 26.39 ± 1.86 | 0.51 | 0.83 | 0.62 |
|  | Injured | 26.59 ± 1.08 | 25.72 ± 1.39 | 25.28 ± 2.60 |  |  |  |
| IL-2 | Control | 40.10 ± 1.50 | 40.93 ± 1.12 | 40.26 ± 1.87 | 0.41 | 0.44 | 0.56 |
|  | Injured | 39.17 ± 2.25 | 40.16 ± 1.20 | 40.98 ± 1.38 |  |  |  |
| IL-3 | Control | 5.07 ± 0.12 | 4.55 ± 0.25 | 5.06 ± 0.39 | 0.29 | 0.45 | 0.67 |
|  | Injured | 4.84 ± 0.35 | 4.83 ± 0.76 | 4.80 ± 0.48 |  |  |  |
| IL-4 | Control | 2.12 ± 0.12 | 2.14 ± 0.19 | 2.19 ± 0.14 | 0.59 | 0.99 | 0.75 |
|  | Injured | 2.2 ± 0.20 | 2.19 ± 0.11 | 2.12 ± 0.20 |  |  |  |
| IL-5 | Control | 6.47 ± 0.45 | 7.00 ± 0.35 | 6.83 ± 0.31 | 0.18 | 0.31 | 0.16 |
|  | Injured | 6.65 ± 0.30 | 6.63 ± 0.26 | 6.44 ± 0.39 |  |  |  |
| IL-6 | Control | 1271.82 ± 746.28 | 1696.15 ± 576.15 | 1354.88 ± 646.921 | 0.58 | 0.61 | **0.032** |
|  | Injured | 2829.57 ± 2675.78 | 3217.03 ± 2685.95 | 1820.83 ± 741.73 |  |  |  |
| IL-7 | Control | 16.15 ± 1.83 | 15.26 ± 1.40 | 16.03 ± 1.62 | 0.17 | 0.50 | 0.94 |
|  | Injured | 14.89 ± 1.04 | 15.91± 0.69 | 16.74 ± 1.89 |  |  |  |
| IL-9 | Control | 49.88 ± 1.70 | 52.47 ± 3.26 | 49.59 ± 3.09 | 0.98 | **0.030** | 0.75 |
|  | Injured | 49.10 ± 1.18 | 52.32 ± 3.18 | 49.30 ± 3.25 |  |  |  |
| IL-10 | Control | 14.01 ± 1.79 | 15.15 ± 1.55 | 14.83 ± 1.48 | 0.50 | 0.61 | 0.54 |
|  | Injured | 14.32 ± 1.59 | 13.60 ± 1.21 | 14.80 ± 1.88 |  |  |  |
| IL-12 (p40) | Control | 18.74 ± 1.95 | 19.84 ± 2.09 | 18.16 ± 2.39 | 0.29 | 0.35 | 0.99 |
|  | Injured | 17.67 ± 1.39 | 19.18 ± 1.66 | 19.87 ± 1.57 |  |  |  |
| IL-12 (p70) | Control | 78.83 ± 4.07 | 80.79 ± 7.88 | 80.05 ± 5.50 | 0.57 | 0.42 | 0.99 |
|  | Injured | 80.93 ± 4.47 | 77.07 ± 3.27 | 81.79 ± 4.25 |  |  |  |
| IL-13 | Control | 5.48 ± 0.55 | 5.23 ± 0.45 | 5.48 ± 0.39 | 0.22 | 0.86 | 0.93 |
|  | Injured | 5.34 ± 0.28 | 5.70 ± 0.55 | 5.19 ± 0.67 |  |  |  |
| IL-15 | Control | 52.71 ± 5.57 | 63.43 ± 7.55 | 58.33 ± 5.58 | **0.031** | 0.38 | 0.41 |
|  | Injured | 62.78 ± 1.12 | 58.49 ± 3.43 | 58.27 ± 4.69 |  |  |  |
| IL-17 | Control | 2.01 ± 0.12 | 2.08 ± 0.13 | 2.05 ± 0.12 | 0.58 | 0.68 | 0.13 |
|  | Injured | 2.12 ± 0.21 | 2.09 ± 0.13 | 2.20 ± 0.16 |  |  |  |
| CXCL10 | Control | 43.74 ± 35.72 | 69.63 ± 28.56 | 68.47 ± 31.92 | 0.62 | 0.11 | 0.58 |
|  | Injured | 42.70 ± 18.33 | 68.66 ± 29.18 | 90.20 ± 37.94 |  |  |  |
| CXCL1 | Control | 2200.74 ± 711.84 | 2152.18 ± 348.92 | 1826.09 ± 314.98 | 0.89 | 0.31 | 0.12 |
|  | Injured | 2365.88 ± 624.20 | 2429.41 ± 436.29 | 1979.68 ± 403.56 |  |  |  |
| LIF | Control | 25.04 ± 8.09 | 31.65 ± 8.76 | 28.99 ± 11.31 | 0.83 | 0.52 | **0.041** |
|  | Injured | 33.64 ± 19.70 | 39.88 ±14.77 | 43.09 ± 9.86 |  |  |  |
| CXCL5 | Control | 121.91 ± 159.40 | 69.74 ± 34.86 | 42.48 ± 17.28 | 0.75 | 0.17 | 0.79 |
|  | Injured | 127.26 ± 82.67 | 93.93 ± 60.87 | 30.49 ± 24.06 |  |  |  |
| CCL2 | Control | 3873.37 ± 1444.87 | 4168.00 ± 1086.71 | 4119.91 ± 741.22 | 0.54 | 0.79 | 0.31 |
|  | Injured | 4480.43 ± 947.33 | 4617.40 ± 791.76 | 3977.46 ± 620.99 |  |  |  |
| MCSF | Control | 20.41 ± 1.82 | 21.24 ± 1.69 | 20.43 ± 1.76 | 0.22 | 0.46 | 0.66 |
|  | Injured | 20.60 ± 2.20 | 20.11 ± 1.20 | 22.30 ± 1.35 |  |  |  |
| CXCL9 | Control | 8.26 ± 3.73 | 13.71 ± 4.60 | 14.99 ± 6.29 | 0.26 | 0.079 | 0.43 |
|  | Injured | 9.52 ± 2.43 | 11.77 ± 3.26 | 19.38 ± 9.51 |  |  |  |
| CCL3 | Control | 107.76 ± 60.80 | 163.98 ± 51.15 | 114.24 ± 35.51 | 0.73 | 0.20 | 0.15 |
|  | Injured | 169.84 ± 103.02 | 187.80 ± 57.58 | 133.57 ± 46.38 |  |  |  |
| CCL4 | Control | 198.04 ± 142.11 | 309.55 ± 129.75 | 209.41 ± 79.26 | 0.72 | 0.34 | **0.031** |
|  | Injured | 359.43 ± 230.50 | 412.91 ± 165.88 | 278.70 ± 127.01 |  |  |  |
| CXCL2 | Control | 431.41 ± 34.85 | 460.03 ± 49.88 | 424.97 ± 21.33 | 0.79 | 0.62 | **0.015** |
|  | Injured | 521.54 ± 144.02 | 520.06 ± 95.10 | 475.79 ± 63.92 |  |  |  |
| CCL5 | Control | 25.35 ± 25.81 | 42.81 ± 21.25 | 20.95 ± 20.97 | 0.90 | 0.062 | 0.53 |
|  | Injured | 30.08 ± 20.43 | 51.06 ± 21.31 | 21.46 ± 11.14 |  |  |  |
| TNFα | Control | 8.30 ± 0.25 | 8.90 ± 0.28 | 8.73 ± 0.14 | 0.054 | 0.71 | 0.065 |
|  | Injured | 9.15 ± 0.38 | 8.69 ± 0.58 | 9.02 ± 0.53 |  |  |  |
| VEGFA | Control | 326.07 ± 80.78 | 256.65 ± 76.07 | 297.34 ± 67.73 | 0.98 | 0.37 | 0.51 |
|  | Injured | 344.27 ± 124.44 | 280.11 ± 49.32 | 309.26 ± 96.02 |  |  |  |
| CCL21 | Control | 130.57 ± 70.48 | 132.46 ± 67.73 | 217.44 ± 175.56 | 0.43 | **0.028** | **0.049** |
|  | Injured | 215.33 ± 16.93 | 155.51 ± 108.44 | 360.02 ± 85.66 |  |  |  |
| CX3CL1 | Control | 334.68 ± 104.26 | 334.20 ± 35.55 | 361.33 ± 49.05 | 0.81 | 0.65 | 0.12 |
|  | Injured | 371.01 ± 63.45 | 347.75 ± 67.32 | 398.35 ± 100.75 |  |  |  |
| IFN- β1 | Control | 32.52 ± 2.50 | 32.46 ± 2.66 | 35.42 ± 1.81 | 0.18 | 0.33 | 0.63 |
|  | Injured | 31.70 ± 3.53 | 35.19 ± 5.04 | 31.39 ± 2.20 |  |  |  |
| IL-11 | Control | 1378.51 ± 792.09 | 817.59 ± 288.39 | 1078.30 ± 558.52 | 0.65 | 0.45 | 0.68 |
|  | Injured | 1019.21 ± 680.27 | 931.61 ± 553.19 | 1061.97 ± 232.88 |  |  |  |
| IL-16 | Control | 52.95 ± 21.55 | 36.79 ± 6.51 | 45.89 ± 2.97 | 0.30 | 0.070 | 0.68 |
|  | Injured | 40.20 ± 13.45 | 39.10 ± 6.15 | 50.57 ± 7.95 |  |  |  |
| IL-20 | Control | OOR< | 1.72^^^ | 2.74 ± 1.44^^^ |  | | |
|  | Injured | OOR< | 4.10 ± 1.80^^^ | OOR< |  |  |  |
| CCL12 | Control | 415.60 ± 264.92 | 782.40 ± 378.12 | 622.77 ± 239.44 | 0.27 | 0.95 | **0.014** |
|  | Injured | 1389.04 ± 1118.43 | 995.51 ± 424.39 | 1016.30 ± 553.30 |  |  |  |
| CCL22 | Control | 15.41 ± 11.83 | 17.08 ± 12.14 | 10.73 ± 2.75 | **0.043** | 0.24 | **0.0038** |
|  | Injured | 45.45 ± 35.19 | 24.68 ± 14.45 | 14.79 ± 6.50 |  |  |  |
| CCL20 | Control | 0.44 ± 0.31 | 0.95 ± 0.44 | 0.44 ± 0.28 | 0.14 | 0.092 | **0.010** |
|  | Injured | 3.04 ± 2.78 | 3.41 ± 2.64 | 0.46 ± 0.14 |  |  |  |
| CCL19 | Control | 22.19 ± 16.20 | 19.48 ± 4.60 | 37.65 ± 15.59 | 0.68 | 0.072 | 0.88 |
|  | Injured | 23.73 ± 8.17 | 23.20 ± 11.66 | 33.99 ± 9.79 |  |  |  |
| CC17 | Control | 12.13 ± 18.65 | 4.10 ± 0.68 | 3.70 ± 1.72 | **0.021** | 0.19 | **0.0083** |
|  | Injured | 28.74 ± 32.78 | 6.67 ± 3.00 | 4.32 ± 1.18 |  |  |  |
| TIMP-1 | Control | OOR> | 140097.31 ± 50267.05^^^ | OOR> |  | | |
|  | Injured | OOR> | 43162.34 ± 29899.94^^^ | OOR> |  |  |  |
| MMP-2 | Control | 83475.00 ± 35374.50 | 86019.39 ± 16572.67 | 80282.12 ± 12693.40 | 0.34 | 0.97 | **0.021** |
|  | Injured | 88882.11 ± 27533.80 | 92545.06 ± 15622.53 | 98050.23 ± 13437.19 |  |  |  |
| MMP-3 | Control | 22520.33 ± 1401.90 | 22330.13 ± 747.91 | 23097.12 ± 1216.39 | 0.20 | 0.62 | 0.061 |
|  | Injured | 24719.01 ± 2939.92 | 23842.42 ± 718.91 | 22792.08 ± 1058.3 |  |  |  |
| MMP-8 | Control | 4396.13 ± 4440.96 | 1632 ± 1087.59 | 1588.46 ± 471.16 | 0.39 | 0.12 | 0.72 |
|  | Injured | 2716.47 ± 977.32 | 2163.85 ± 332.63 | 1969.47 ± 877.49 |  |  |  |
| proMMP-9 | Control | 15359.94 ± 11683.98 | 6358.95 ± 3911.44 | 7590.48 ± 3426.23 | 0.13 | 0.25 | 0.69 |
|  | Injured | 7821.78 ± 3828.74 | 6476.04 ± 2996.21 | 12064.76 ± 6720.95 |  |  |  |
| MMP-1 | Control | 34082.41 ± 28399.01 | 26406.51± 21341.94 | 14846.65 ± 8004.04 | 0.41 | 0.34 | 0.092 |
|  | Injured | 64162.40 ± 77011.70 | 32676.60 ± 21773.46 | 21201.40 ± 7406.39 |  |  |  |

OOR<: Out of range below curve

OOR>: Out of range above curve

^^^Subset of samples
