## Supplementary figures and images for "The progression of neurovascular features and chemokine signatures of the intervertebral disc with degeneration"

### {Supplemental}

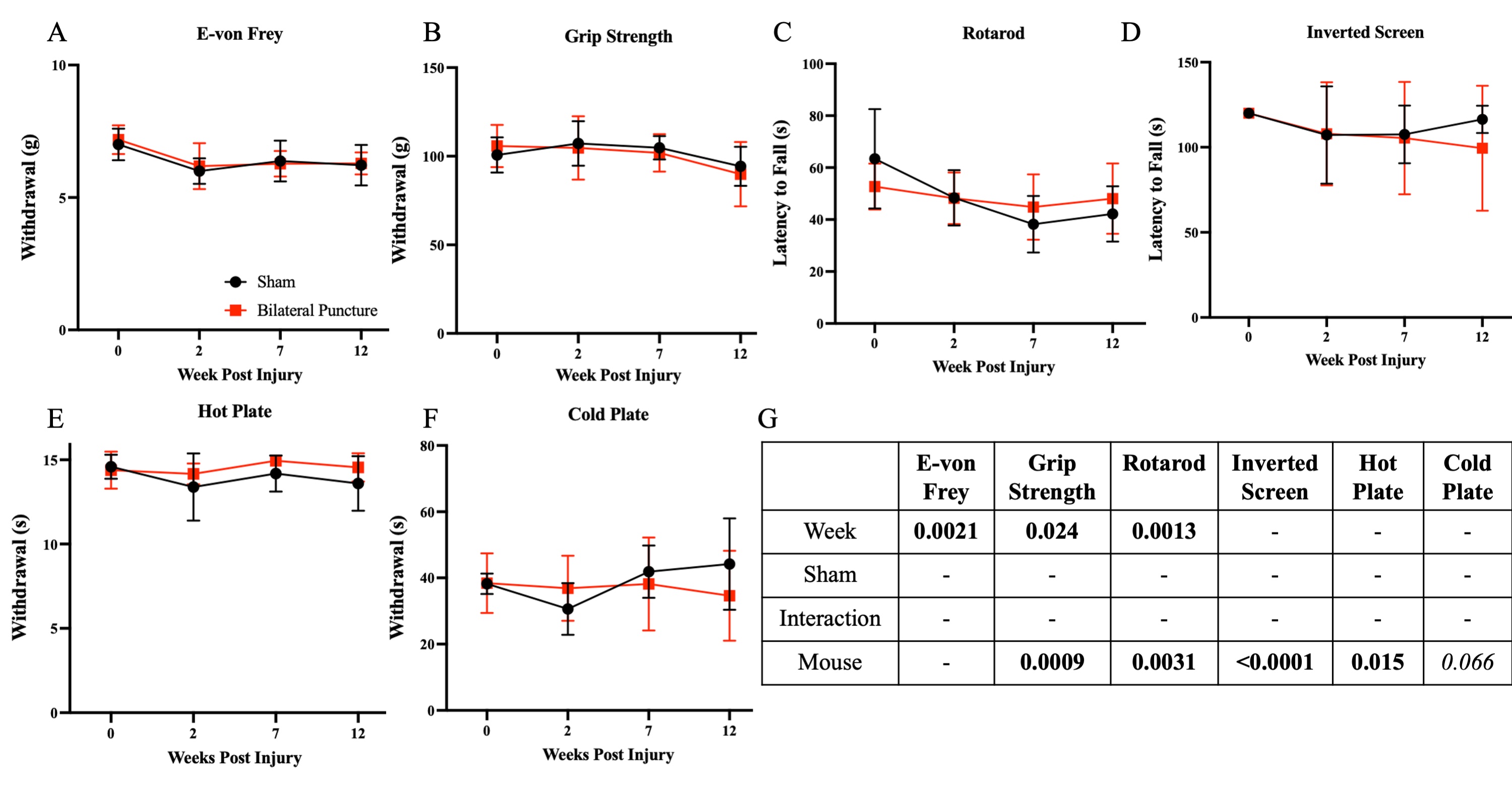

### {Supplemental}

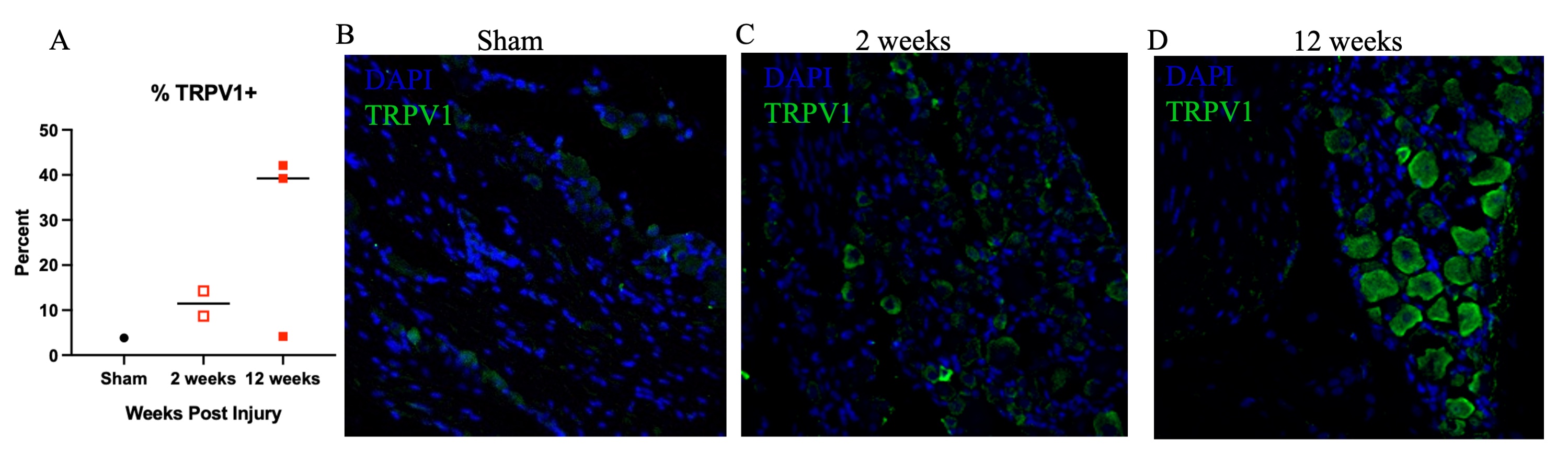
